# Impaired migration and premature differentiation underlie the neurological phenotype associated with PCDH12 loss of function

**DOI:** 10.1101/2023.01.05.522934

**Authors:** Jennifer Rakotomamonjy, Lauren Rylaarsdam, Lucas Fares-Taie, Sean McDermott, Devin Davies, George Yang, Fikayo Fagbemi, Maya Epstein, Alicia Guemez-Gamboa

## Abstract

Protocadherins (PCDHs) are cell adhesion molecules that regulate many essential neurodevelopmental processes related to neuronal maturation, dendritic arbor formation, axon pathfinding, and synaptic plasticity. Bi-allelic loss-of-function variants in *PCDH12* are associated with several neurodevelopmental disorders (NDDs) such as diencephalic-mesencephalic dysplasia syndrome, cerebral palsy, cerebellar ataxia, and microcephaly. Despite the highly deleterious outcome resulting from loss of PCDH12, little is known about its role during brain development and disease. Here, we show that PCDH12 loss severely impairs cerebral organoid development with reduced proliferative areas and disrupted laminar organization. 2D models further show that neural progenitor cells lacking PCDH12 prematurely exit cell cycle and differentiate earlier when compared to wildtype. Furthermore, we show that PCDH12 regulates neuronal migration through a mechanism requiring ADAM10-mediated ectodomain shedding and membrane recruitment of cytoskeleton regulators. Our data demonstrate a critical and broad involvement of PCDH12 in cortical development, revealing the pathogenic mechanisms underlying PCDH12-related NDDs.

## Introduction

Brain development involves key phases that are tightly regulated to ensure proper function. Successful cortical development involves mainly three crucial steps. The first is generation of the correct cell type at the right time and in the right proportions during neurogenesis. Second, the newborn neuron must reach its proper terminal location, generating the well-characterized organization in six cortical layers. Third, neurons must mature and establish proper connections with the right partners to ensure a functional neuronal network. All three steps heavily depend on transmembrane cell adhesion molecules. These proteins provide a two-sided platform: their extracellular domain mediates cell-cell interaction/recognition and adhesion, while the intracellular tail initiates signal transduction pathways and reorganization of the actin cytoskeleton.

Protocadherins (PCDHs) are cell adhesion molecules that belong to the calcium-dependent cadherin superfamily. Expressed predominantly in the nervous system, PCDHs have been shown to regulate many essential neurodevelopmental processes related to neuronal maturation, dendritic arbor formation, axon pathfinding, and synaptic plasticity (Flaherty and Maniatis, 2020). It is then no surprise that pathogenic variants in *PCDH* genes are associated with various neurodevelopmental disorders (NDDs) such as epilepsy, autism spectrum disorders, and intellectual disability (Anitha et al., 2013; Dibbens et al., 2008; Morrow et al., 2008).

We recently reported that bi-allelic loss-of-function variants in *PCDH12* also result in NDD. PCDH12 is part of the δ2 subgroup of non-clustered PCDHs, which possess six extracellular cadherin (EC) domains that mediate homophilic interactions. The PCDH12 intracellular tail contains the WAVE Regulatory Complex (WRC) interacting receptor sequence (WIRS) that regulates actin polymerization (Chen et al., 2014). PCDH12 loss of function results in a wide neurological phenotypic spectrum including cerebral palsy, microcephaly, intellectual disability, speech impairment, seizures, and psychomotor delays. Brain imaging further revealed brainstem malformations, white matter abnormalities, cerebellar ataxia, and brain calcifications (Aran et al., 2016; Fazeli et al., 2022; Guemez-Gamboa et al., 2018; Mattioli et al., 2021; Suzuki-Muromoto et al., 2018; Vineeth et al., 2019). Despite such severe neurodevelopmental abnormalities resulting from PCDH12 deficiency, little is known about its function in brain development and disease.

To decipher the pathogenic mechanisms leading to PCDH12-related NDDs, we generated *PCDH12* knockout (*PCDH12*-KO) human stem cell lines and investigated the effect of PCDH12 loss in cerebral organoids, neural progenitor cells, and neurons. Our findings show that PCDH12 is essential for proper development of cerebral organoids, playing a critical role in neural progenitor (NPC) differentiation and establishment of correct progenitor and neuron layers. We also found evidence that PCDH12 affects neuronal migration through a mechanism involving metalloproteinase-mediated ectodomain shedding. Altogether, these findings illuminate the critical role of PCDH12 during human cortical development and establish a novel mechanism for PCDH12-related NDDs.

## Results

### Loss of PCDH12 impairs cerebral organoid morphology, ventricular-like zone thickness, and laminar organization

Prior studies have reported that *Pcdh12*-knockout mice are viable and fertile, with no major cortical abnormalities or morphological changes compared to wild type mice (Rampon et al., 2005). Thus, to study the mechanisms by which loss of PCDH12 affects brain development, we turned to CRISPR/Cas9 gene editing to insert frameshift variants into exon 1 to lead to complete PCDH12 loss (*PCDH12*-KO) in human stem cells (Cong et al., 2013; Ran et al., 2013). Moreover, we also inserted a specific frameshift causing variant (c.2511delG) that has been reported in multiple patients **(Supplemental Figure S1 and S2A)**. We next generated 3D cerebral organoid cultures (Lancaster and Knoblich, 2014; Lancaster et al., 2013) **(Figure 1A, Supplemental Figure S2B and C)**. Morphological differences became apparent between *PCDH12*-KO organoids and wild-type (WT) controls very early during the differentiation process. On day 7, proper neuroectodermal differentiation is indicated by a round embryoid body (EB) with bright and smooth edges. Though optically clear edges were present in *PCDH12*-KO EBs, they lost their circular shape compared to WT EBs (**Figure 1B and 1C**). By day 10, matrix-embedded organoids should develop neuroepithelial buds at the organoid surface, resulting in decreased circularity. WT organoids properly showed a 1.2-fold decrease in circularity between day 7 and day 10 compared to a 1.05-fold decrease for *PCDH12*-KO organoids (**Figure 1C**). Because organoid differentiation is an inherently heterogenous process, we used a scoring system to rate successful differentiation at day 10, from category 1 showing ideal organoids that have well developed neuroepithelial buds and optically clear edges, to category 4 representing failed organoids that lack surface budding and optically clear edges. *PCDH12*-KO organoids presented with reduced epithelial expansion with 88% of them belonging to impaired categories 3 and 4, while WT cultures generated category-1 and -2 organoids only (**Figure 1D and 1E**).

**Figure 1:**
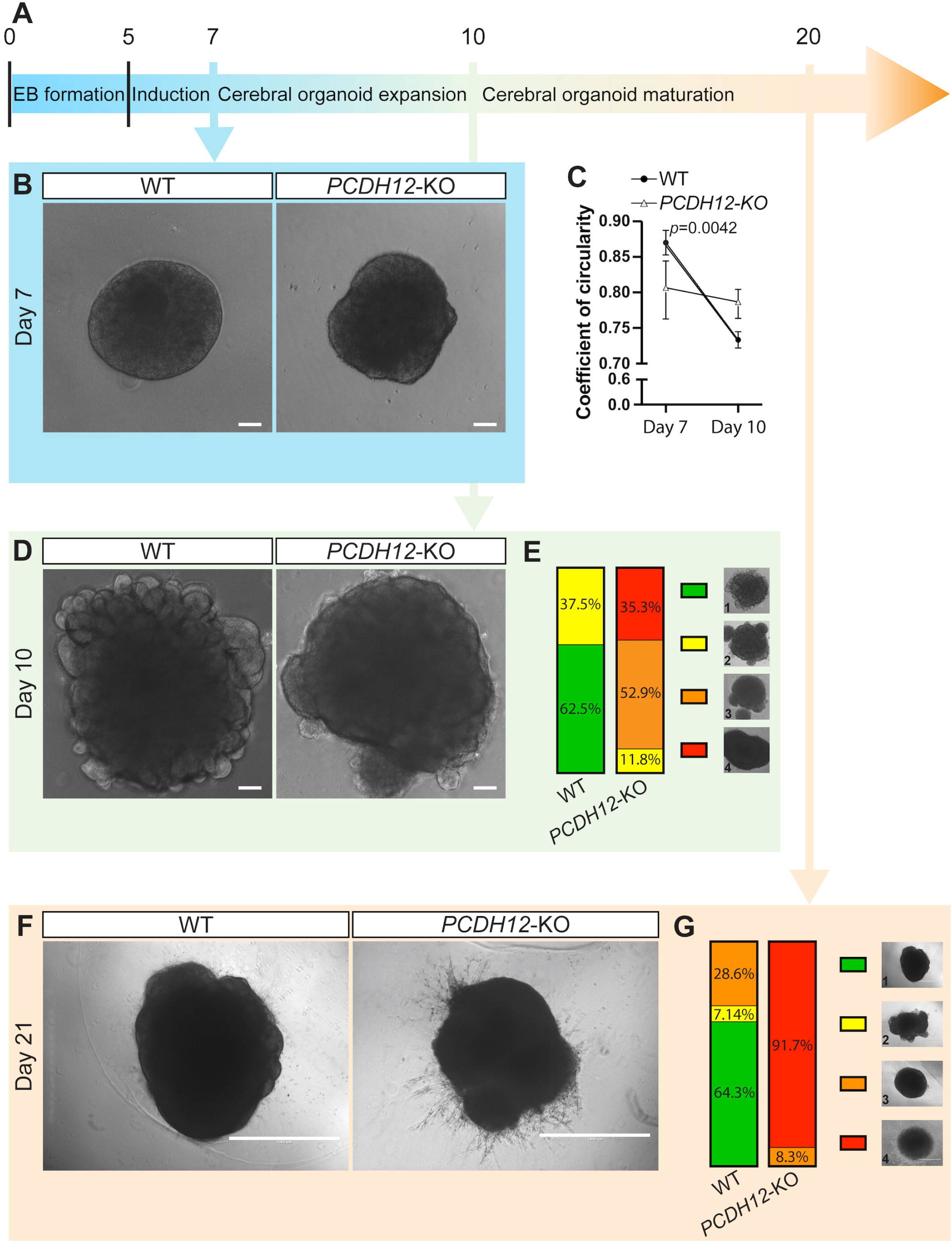
Loss of PCHD12 impairs cerebral organoid morphology. (A) Schematic diagram of the timeline of the cerebral organoid formation. (B) Representative bright-field images of WT (left) and *PCDH12*-KO (right) embryoid bodies (EBs) at day 7, before Matrigel embedding, showing a brightening of organoid edges, indicative of ectodermal differentiation. (C) Quantification of EB coefficient of circularity, defined as 4π(area/perimeter^2^) at day 7 (n=38 for WT; n=39 for PCDH12-KO; mean ± SD) and day 10 (n=32 for WT; n=34 for PCDH12-KO; mean ± SD) *p*=0.0042; Comparison of Fits. (D) Representative bright-field images WT (left) and *PCDH12*-KO (right) EBs at day 10, showing more pronounced surface budding, indicative of neuroepithelial expansion, in WT compared to *PCDH12*-KO EBs. (E) Percentages of EBs in developmental categories 1 to 4 (right). Category 1: Uniform and pronounced surface budding, optically clear edges, as shown in D, left panel; Category 2: smoother surface budding, with occasional non-neuroepithelial outgrowths, as shown in C, right pfanel; Category 3: few surface budding with recurrent non-neuroepithelial outgrowths; Category 4: No surface budding, lack of optically clear edges. (F) Representative category-1 WT (left) and category-3 *PCDH12*-KO (right) cerebral organoids at day 21. (G) Percentages of cerebral organoids in developmental categories 1 to 4. Category 1: Expanded cerebral tissue with ventricle-like structures, as shown in F, left panel; Category 2: Expanded cerebral tissue with ventricle-like structures, alongside non-neuronal outgrowths; Category 3: Minimal cerebral expansion with fewer and smaller ventricle-like structures. Category 4: No cerebral expansion, with cell processes extending into the Matrigel. Scale bars represent 100 μm (B and D) and 1 mm (F).

By day 21, expanded cerebral tissue with ventricle-like structures are expected, as observed in WT organoids (**Figure 1F**). In contrast, most *PCDH12*-KO organoids showed little-to-no cerebral expansion and multiple cell processes extending into the embedding matrix, suggesting premature neural differentiation (Lancaster and Knoblich, 2014) (**Figure 1F and 1G**). Moreover, day 21 *PCDH12*-KO organoids showed a reduced number of N-cadherin polarized neural rosettes per organoid when compared to WT (**Figure 2A and 2B**). Neural rosettes lacking PCDH12 also displayed a reduced number of apically anchored mitotic cells (**Figure 2C and 2D**). This points to a deficient proliferative area, as evidenced by the decrease of the ventricular-like zone thickness defined by SOX2+ progenitors (**Figure 2E and 2F**).

**Figure 2:**
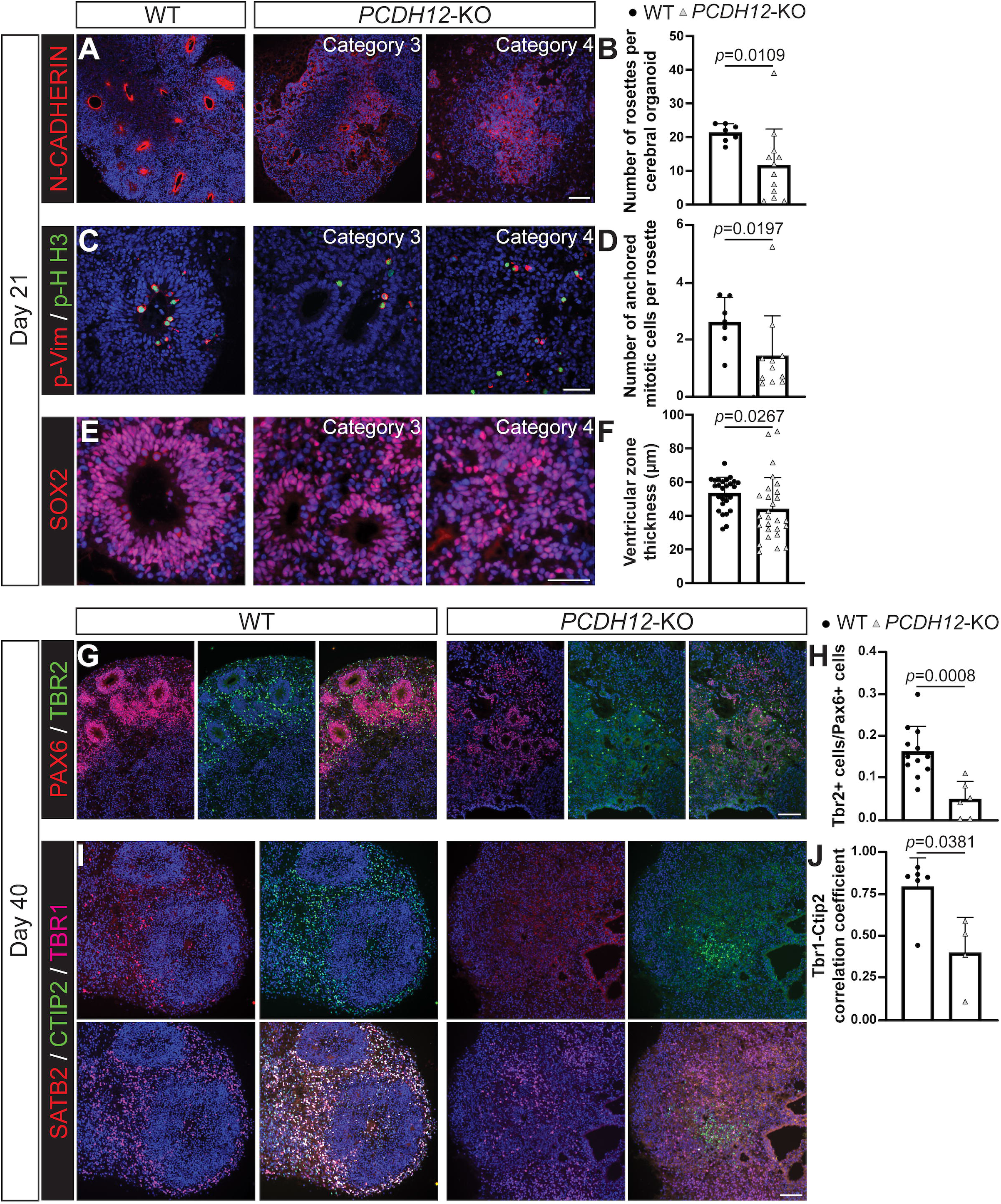
Reduced ventricular zone and impaired laminar organization in *PCDH12*-KO cerebral organoids. (A) Immunofluorescence images of WT (left), category 3 (middle), and category 4 (right) *PCDH12*-KO cerebral organoids at day 21 stained with N-cadherin (red). (B) Quantification of N-cadherin-defined cortical rosettes in day-21 cerebral organoids. Each point represents the number of rosettes for one cerebral organoid (n=7 organoids for WT; n=12 organoids for *PCDH12*-KO; mean + SD; Unpaired t test with Welch’s correction). (C) Immunofluorescence images of WT (left), category 3 (middle), and category 4 (right) *PCDH12*-KO cerebral organoids at day 21 stained with phospho-vimentin (p-Vim, red), and phospho-Histone H3 (pH-H3, green). (D) Percentages of cortical rosettes carrying anchored mitotic cells in day-21 cerebral organoids. Each point represents the percentage for one cerebral organoid (n=6 organoids for WT; n=6 organoids for *PCDH12*-KO; mean + SD; Unpaired t test with Welch’s correction). (E) Immunofluorescence images of WT (left), category 3 (middle), and category 4 (right) *PCDH12*-KO cerebral organoids at day 21 stained with Sox2 (red). (F) Quantification of ventricular zone thickness in day-21 cerebral organoids, defined by the thickness of the layer of SOX2+ cells. Each point represents the average ventricular zone thickness for one cortical rosette (n=27 rosettes for WT; n=27 rosettes for *PCDH12*-KO; mean + SD; Unpaired t test with Welch’s correction). (G) Immunofluorescence images of WT (3 left panels) and *PCDH12*-KO (3 right panels) cerebral organoids at day 40 stained with Pax6 (red) and Tbr2 (green). (H) Average ratio of Tbr2-relative to Pax6-positive cells per rosette in day 40 organoids (n=12 rosettes for WT; n=6 rosettes for *PCDH12*-KO; mean + SD; Unpaired t test). (I) Immunofluorescence images of WT (4 left panels) and *PCDH12*-KO (4 right panels) cerebral organoids at day 40 stained with Satb2 (red), Ctip2 (green), and Tbr1 (magenta). (J) Average Ctip2/Tbr1 Pearson’s correlation coefficients showing degree of colocalization in Day 40 organoids (n=6 organoids for WT; n=4 organoids for *PCDH12*-KO; mean + SD; Mann-Whitney test). Scale bars represent 100 μm (A, G, and I) and 50 μm (C and E).

By day 40, the neuroprogenitor zone was further reduced in *PCDH12*-KO organoids, displaying rosettes with very thin PAX6+ radial glial cell (RGC) layers compared to WT rosettes (**Figure 2G**). At this stage, RGCs can follow two paths: 1) the indirect neurogenesis route, consisting of asymmetrical divisions to generate another PAX6+ daughter cell and a TBR2+ intermediate progenitor (Englund et al., 2005) that will in turn divide to generate neurons; or 2) the direct neurogenesis, skipping the intermediate progenitor generation and directly differentiating to neurons (Haubensak et al., 2004; Miyata et al., 2004; Noctor et al., 2004). Our results indicate that loss of PCDH12 leads to a decreased proportion of TBR2+ intermediate progenitors, indicating a propensity towards direct neurogenesis compared to WT (**Figure 2H**). Past the progenitor state, neurons lacking PCH12 failed to organize in a basal layer around the progenitor layer (**Figure 2I**). Furthermore, TBR1+ and CTIP2+ early-born neurons strongly colocalized in WT organoids at day 40, indicating fate specification onset of postmitotic neurons. However, segregation of TBR1 and CTIP2 expression was well advanced in *PCDH12*-KO organoids, as evidenced by a 50% decrease in the TBR1-CTIP2 signal correlation coefficient (**Figure 2J**), suggesting that neuronal differentiation proceeds prematurely and/or more rapidly in absence of PCDH12. Taken together, our results suggest that PCDH12 is involved in the maintenance of the progenitor state and the proper lamination of cortical neurons.

### PCDH12 loss results in compromised cell cycle re-entry and premature neuronal differentiation

Because we observed accelerated neuronal differentiation and aberrant lamination in *PCDH12*-KO organoids, we speculated that several developmental events may be disrupted when PCDH12 is lacking, including proliferative and/or differentiation defects as well as impaired migration preventing proper lamination. To dissect those mechanisms independently and circumvent the developmental variability inherent to the organoid culture process, we generated 2D cultures of neural progenitor cells (2D-NPCs) from our *PCDH12*-KO human stem cell lines (Chambers et al., 2009). We first assessed the proliferative potential of WT and *PCDH12*-KO 2D-NPCs by carrying BrdU pulse-chase experiments. Both short and long pulses did not yield significant differences between genotypes, with both lines showing similar rates of BrdU intake, proportions of actively cycling cells, and comparable cycling speed (**Figure 3A**). However, we observed a 21% decrease in *PCDH12*-KO 2D-NPC cell cycle re-entry at 24 hours when compared to WT, suggesting early exit from the progenitor state (**Figure 3B**).

**Figure 3:**
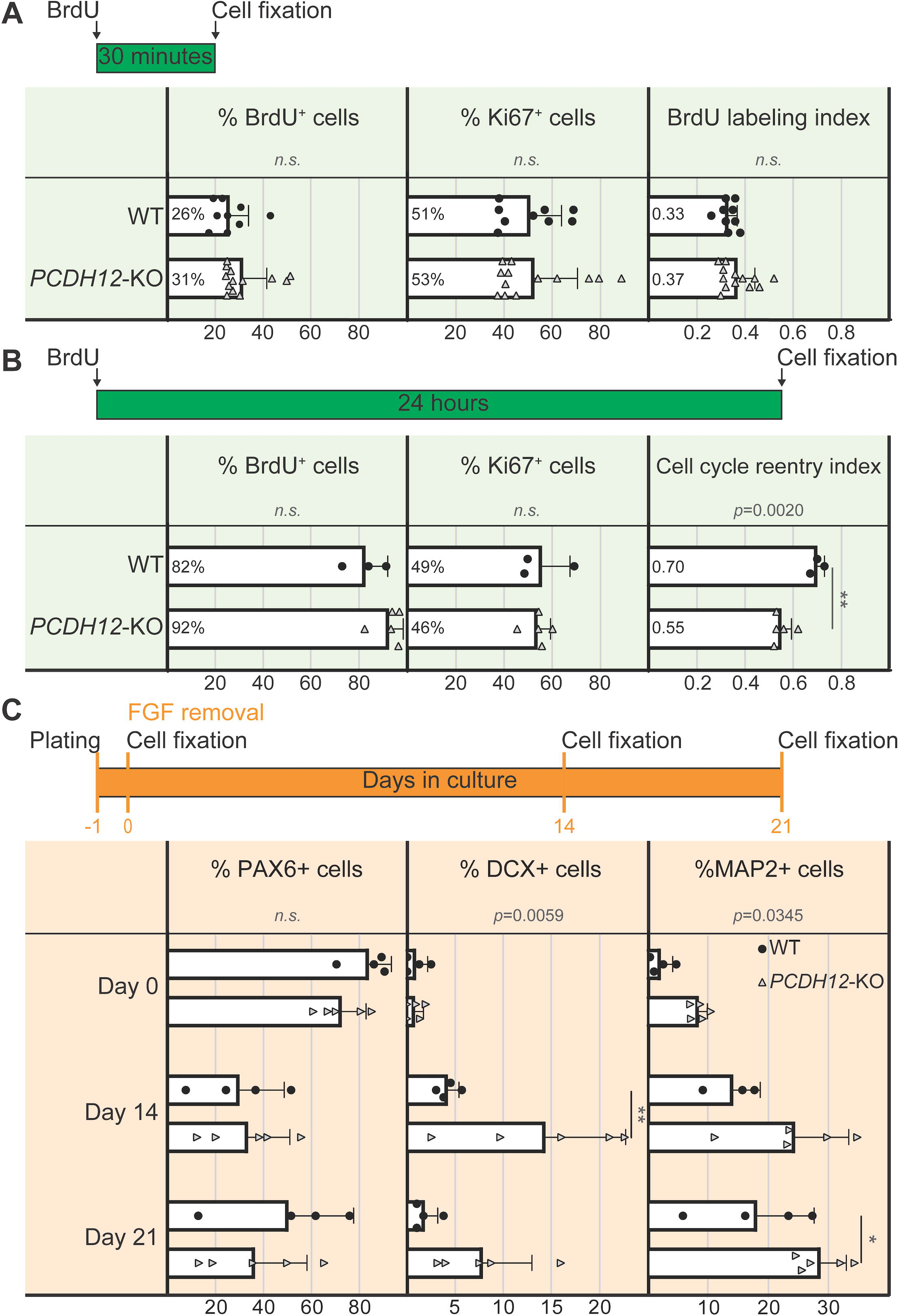
*PCDH12*-KO neural progenitor cells show premature cell cycle withdrawal and early neuronal differentiation. (A) Quantifications of the percentages of BrdU+ and Ki67+ cells, and BrdU labeling index defined as double-positive BrdU+ - Ki67+ cells/total Ki67+ cells after a short 30-minute BrdU pulse. Each data point represents one biological replicate (n=9 for WT; n=13 for *PCDH12*-KO; mean + SD; Unpaired t test). (B) Quantifications of the percentages of BrdU+ and Ki67+ cells, and cell cycle re-entry index defined as double-positive BrdU+ - Ki67+ cells/BrdU+ cells after a long 24-hour BrdU pulse. Each data point represents one biological replicate (n=3 for WT; n=5 for *PCDH12*-KO; mean + SD; Mann-Whitney test). (C) Percentages of PAX6+, doublecortin (DCX)+, and MAP2+ cells at the removal of FGF (Day 0), 14 days, and 21 days post-FGF removal. Each data point represents one biological replicate (n=4 for Pax6 and DCX for WT at day 0, 14, and 21; n=5 for PAX6, DCX, and MAP2 for *PCDH12*-KO at day 0, 14, and 21; n=4 at day 0 and 21, and n=3 at day 14 for MAP2 for WT; mean + SD; Multiple unpaired t tests).

We next investigated the possible outcome for these cells prematurely exiting the cell cycle. No change in cell death rate as measured by TUNEL and caspase-3 activity was observed (**Supplemental Figure S3**). We then induced neuronal differentiation by withdrawing fibroblast growth factor (FGF) from the cell culture medium. As expected, the proportion of PAX6+ cells kept decreasing over time in both WT and KO 2D-NPC cultures. However, *PCDH12*-KO cells showed increased generation of immature doublecortin+ (DCX+) neurons and mature MAP2+ neurons 14 and 21 days after FGF removal respectively, when compared to WT (**Figure 3C**). Altogether, our results from 2D-NPC cultures support what we observed in organoids: the loss of PCDH12 leads to premature exit from the cell cycle and subsequent early neuronal differentiation.

### *PCDH12*-KO neurons display migration defects

Our cerebral organoid system also revealed that *PCDH12*-KO lines were unable to form proper laminated patterning of developing neural cells (**Figure 2I**). As neuronal migration is crucial for correct cortical layering, we next assessed the migration capabilities of neurons lacking PCDH12. We performed a neurosphere migration assay modified from previously described protocols (Youn et al., 2009) (**Figure 4A**). We found no differences in the distribution of neurosphere-derived migrating neurons between genotypes 48 hours post-plating (**Figure 4B and 4C**). After five days, 56% of *PCDH12*-KO neurons accumulated within the first 100 μm around the neurosphere border versus 38% in WT. On the other hand, only 26% of KO neurons traveled beyond 150 μm versus 45% in WT (**Figure 4D and 4E**), suggesting neural migration defects in neurons lacking PCDH12. To investigate why *PCDH12*-KO neurons do not travel as far as WT neurons, we tracked neuronal migration for 16 hours (**Figure 4F, Supplemental movies S1 and S2**). We found that in the absence of PCDH12, neurons do not maintain directional persistence (**Figure 4G and 4H**), resulting in a 2.9-fold decrease in the average Euclidean distance travelled (**Figure 4I**). Together, our results suggest that impaired migration could explain the defective lamination observed in *PCDH12*-KO organoids.

**Figure 4:**
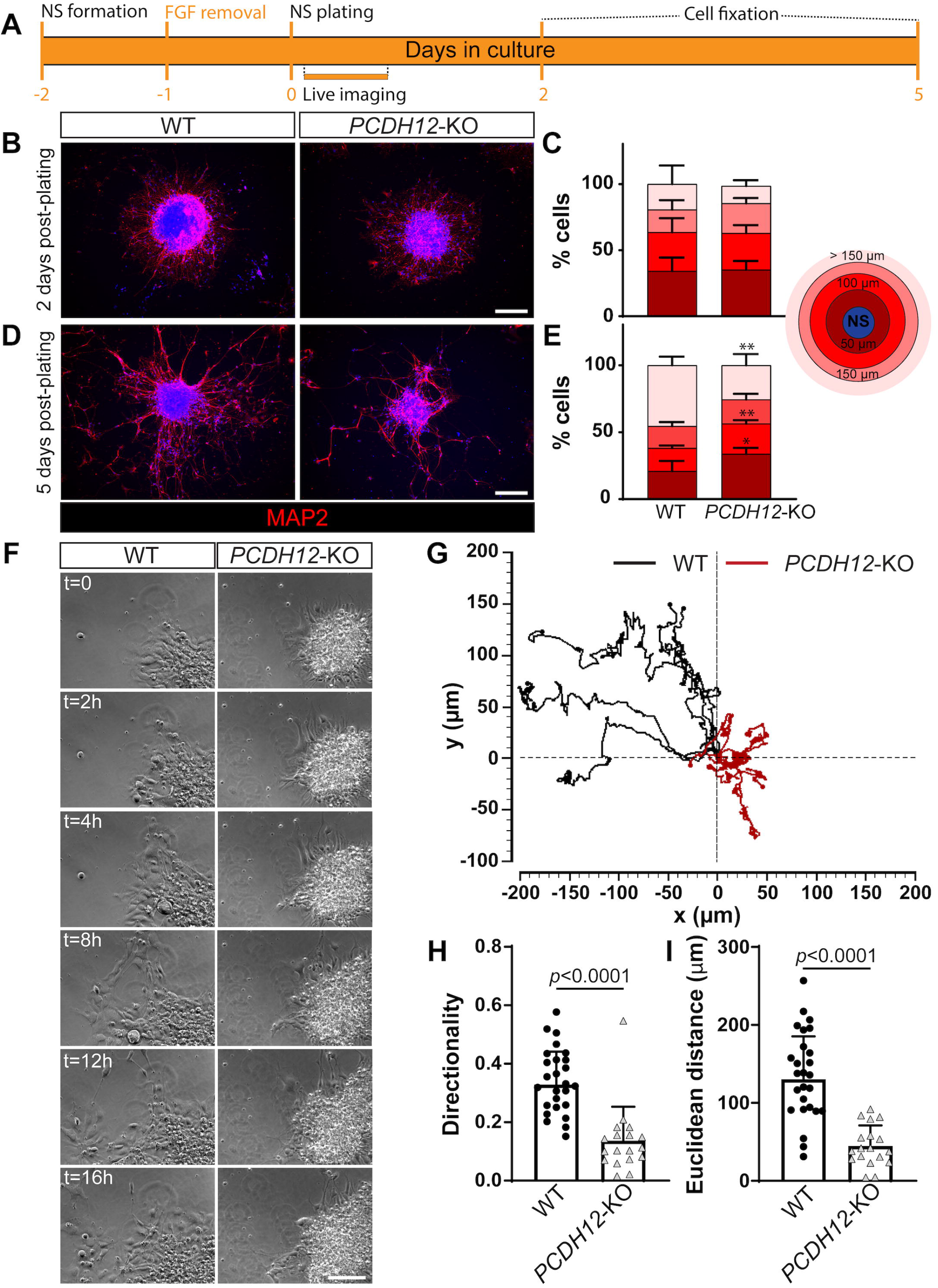
PCDH12 loss of expression leads to impaired neuronal migration. (A) Schematic diagram of the timeline of the neurosphere migration assay. Cell media was half-changed every other day after neurosphere plating. (B) Representative images of WT (left) and *PCDH12*-KO (right) neurospheres stained with MAP2 (red) 48 hours post-plating. (C) Percentages of cells distributed within 50-micron incremental bins from the edge of the neurosphere 48 hours post-plating (n=7 neurospheres for WT; n=16 neurospheres for *PCDH12*-KO; mean + SD; Multiple unpaired t-tests). (D) Representative images of WT (left) and *PCDH12*-KO (right) neurospheres stained with MAP2 (red) 5 days post-plating. (E) Percentages of cells distributed within 50-micron incremental bins from the edge of the neurosphere 5 days post-plating (n=4 neurospheres for WT; n=7 neurospheres for *PCDH12*-KO; mean + SD; ** *p*=0.0067 between WT and *PCDH12*-KO cells in the 50-micron bin; *** *p*=0.0002 between WT and *PCDH12*-KO cells in the bin > 150 microns; Multiple unpaired t-tests). (F) Representative 16-hour live-imaging sequence for WT (left) and *PCDH12*-KO (right) neurospheres. (G) Representative migration plots of five WT (black) and five *PCDH12*-KO (red) migrating neurons over 16 hours. All tracks starting points were normalized to a common origin. (H and I) Quantifications of directionality of neuron motion (H) (a value of 1 translates to a straight-line migration from the starting point) and Euclidean distance (I) (straight line between migration start and end points). Each data point represents a neuron (n=30 neurons for WT; n=18 neurons *PCDH12*-KO; data were acquired from 4 independent experiments; mean + SD; Mann-Whitney test). Scale bars represent 100 μm (B and D) and 200 μm (F).

### PCDH12 function involves recruitment of the WRC to the plasma membrane and regulation of actin polymerization in neural progenitor cells

Neurogenesis and neuronal migration both depend on proper regulation of the actin cytoskeleton. Disruption of cytoskeleton dynamics results in cortical malformations in mice (Cappello et al., 2012) and is observed in patients suffering from NDDs (Griesi-Oliveira et al., 2018; Weitzdoerfer et al., 2002). Actin dynamics are regulated by the WRC, which promotes Arp2/3-mediated actin nucleation (Chen et al., 2010).

Downstream regulators such as cofilin control the balance between polymerization/stabilization and depolymerization/turnover of actin (Ghosh et al., 2004). Since the PCDH12 intracellular domain contains the WIRS motif that binds to the WRC (Chen *et al*., 2014), we tested whether PCDH12 promotes WRC membrane recruitment and subsequent actin polymerization (**Figure 5A**). We observed significantly more WAVE1 - the main component of the WRC - in the plasma membrane fraction from WT 2D-NPCs compared to *PCDH12*-KO cells, while protein levels were similar in cytosolic fractions and whole-cell lysates (**Figure 5B, C, and Supplemental Figure S4A and B**). This result supports that PCDH12 recruits the WRC at the plasma membrane. Decrease in WAVE-1 membrane recruitment did not affect the ability for *PCDH12*-KO neurons to form lamellipodia (**Supplemental Figure S4C**). We further investigated the effect of PCDH12 loss on actin cytoskeleton dynamics by assessing the state of actin polymerization. We found that the ratio of both filamentous polymer (F-actin) to monomeric (G-actin) actin and inactive (phosphorylated) cofilin to active cofilin are increased in *PCDH12*-KO NPCs when compared to WT (**Figure 5E, F**). This indicates that loss of PCDH12 leads to a less dynamic actin cytoskeleton with lower actin turnover.

**Figure 5:**
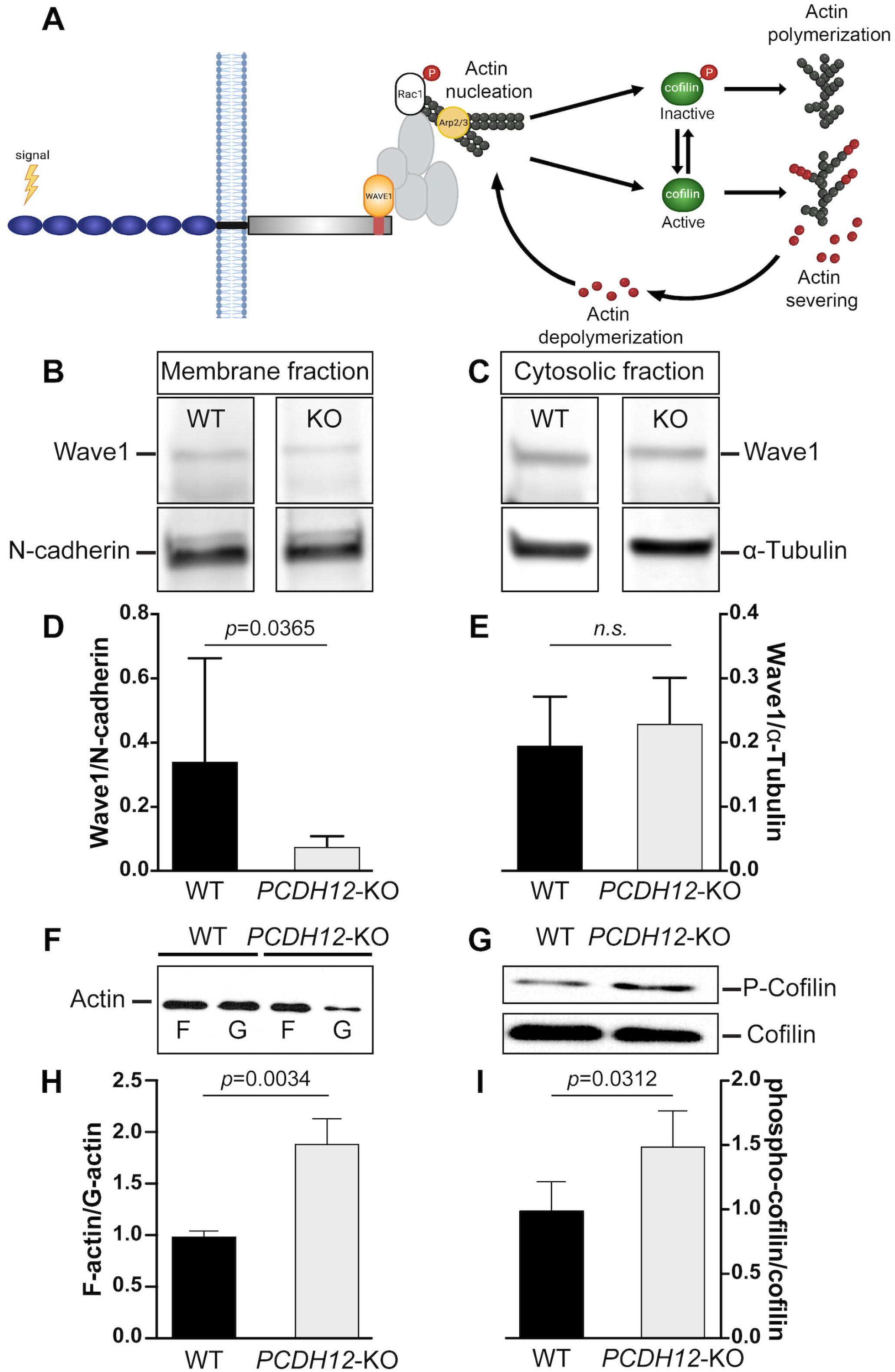
PCDH12 recruits WAVE1 at the plasma membrane and affect actin cytoskeleton regulation. (A) Schematic illustrating the potential interaction between PCDH12 and the actin cytoskeleton. (B and C) Representative western blots of WT and *PCDH12*-KO membrane (B) and cytosolic fractions (C) from neural progenitor cells (NPCs). (D) WAVE1 protein quantification in NPC membrane fractions, relative to N-cadherin (data were acquired from independent experiments; mean + SD; Mann-Whitney test). (E) WAVE1 protein quantification in NPC cytosolic fractions, relative to α -Tubulin (data were acquired from independent experiments; mean + SD; Mann-Whitney test). (F) Representative immunoblots of F- and G-actin in WT and *PCDH12*-KO NPCs. (G) Representative western blot of the presence of total and phospho-cofilin in NPCs. (H) Quantification of the F/G actin ratio (n=3 independent experiments; mean + SD; Unpaired t test). (I) Ratio of phosphorylated to total cofilin in WT and *PCDH12*-KO NPCs (n=4 independent experiments; mean + SD; Unpaired t test).

### Prevention of PCDH12 cleavage reproduces *PCHD12*-KO migration defects

We next asked what upstream signaling event initiated PCDH12-mediated recruitment of the WRC to the plasma membrane. PCDHs are known to favor homophilic *trans* interactions to mediate their function (Harrison et al., 2020). However, direct neuron-neuron contact did not appear to influence direction persistence in our migration assay (**Figure 4**). PCDH12 undergoes cleavage by A Disintegrin And Metalloprotease 10 (ADAM10) to release its extracellular domain and has been shown to affect cell migration *in vitro* (Bouillot et al., 2011). Moreover, ectodomain shedding is an important player in a wide array of biological processes, including brain development (Martin-de-Saavedra et al., 2022). We thus hypothesized that PCDH12 exerts its function through a homophillic-dependent mechanism, with the PCDH ectodomain being its own ligand.

To test this hypothesis, we performed the neurosphere migration assay described earlier in the presence of ADAM10 inhibitory GI 254023X. Our results show that exposing WT neurons to 10 μM GI 254023X led to a disrupted migration pattern similar to what we observed with *PCDH12*-KO neurons. 37% more *PCDH12*-KO neurons remained within the first 50 μM and 43% less migrated beyond 150 μM from the neurosphere edge with the inhibitor treatment compared to control WT cultures (**Figure 6A, B**). We also confirmed that this effect was specific to PCDH12 as we did not observe any significant difference of migratory pattern between control and GI 254023X-treated *PCDH12*-KO neurons (**Figure 6C, D**).

**Figure 6:**
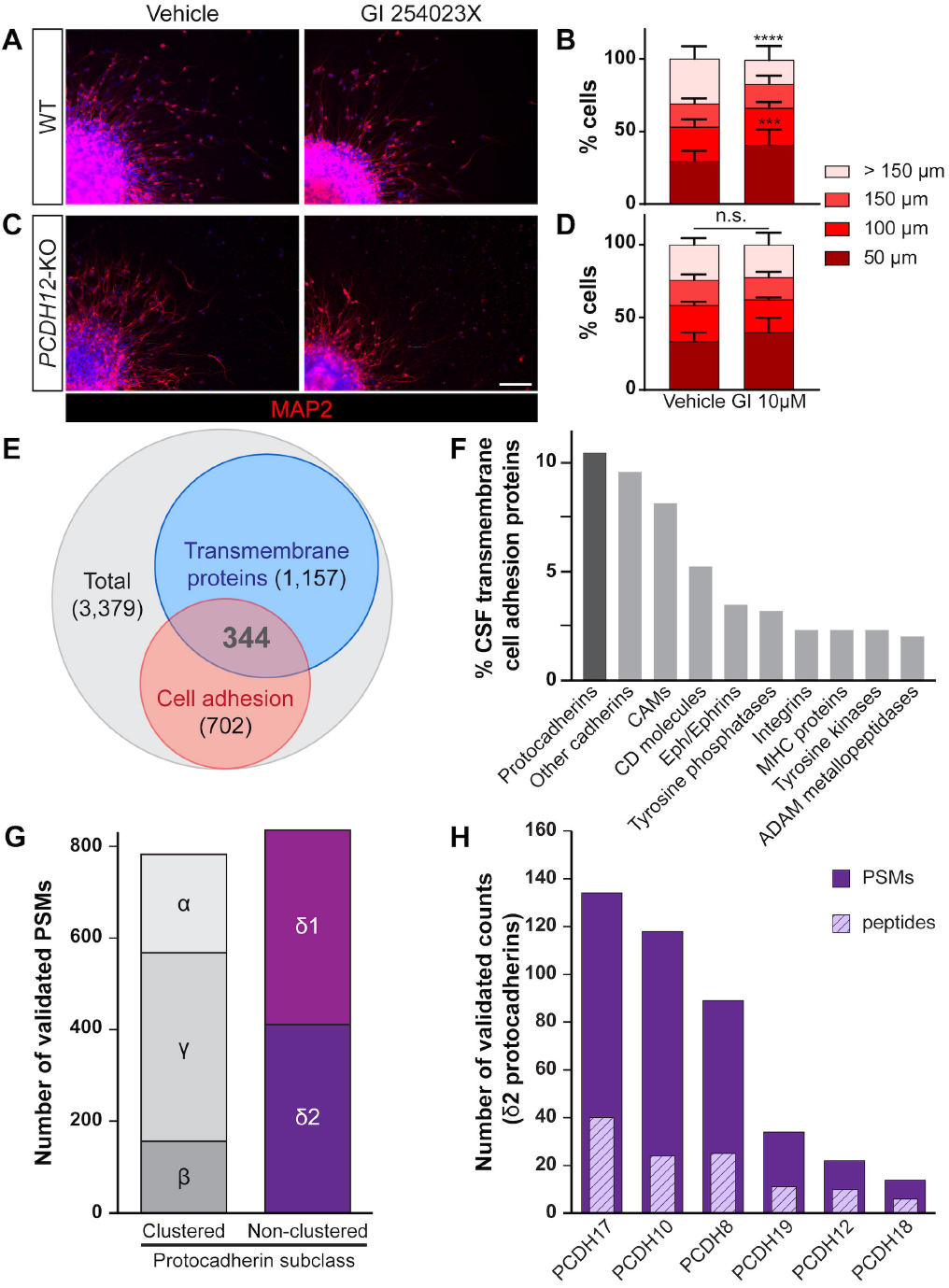
PCDH12 ectodomain is detected in the CSF and is involved in neuronal migration. (A) Representative images of WT neurospheres stained with MAP2 (red) 5 days post-plating treated with vehicle (left) or ADAM-10 inhibitor GI 254023X (right). (B) Average percentages of cells distributed within 50-micron incremental bins from the edge of WT neurospheres (n=15 vehicle-treated neurospheres and n=18 GI 254023X-treated neurospheres; Mean + SD; Brown-Forsythe and Welch ANOVA tests). (C) Representative images of *PCDH12*-KO neurospheres stained with MAP2 (red) 5 days post-plating treated with vehicle (left) or GI 254023X (right). (D) Average percentages of cells distributed within 50-micron incremental bins from the edge of *PCDH12*-KO neurospheres (n=7 vehicle-treated *PCDH12*-KO neurospheres and n=5 GI 254023X-treated *PCDH12*-KO neurospheres; Mean + SD; Brown-Forsythe and Welch ANOVA tests). (E) Venn diagram of total proteome from a commercial pooled CSF sample showing the overlap between transmembrane and cell adhesion proteins (Macron *et al*., 2018). (F) Composition of CSF transmembrane cell adhesion proteins (Macron *et al*., 2018). (G) Total number of peptides for each subclass of protocadherins present in CSF obtained through lumbar puncture (Kroksveen *et al*., 2017). (H) Total number of validated peptide spectrum matches (PSMs) and peptides (hatched bars) among δ2-protocadherins. Scale bar represents 100 μm (A and C).

Our data prompted us to verify the presence of PCDH12 ectodomain in human cerebrospinal fluid (CSF), given that PCDH12 is cleaved *in vivo* and can be detected in human urine and serum (Bouillot *et al*., 2011). We reanalyzed data from previous proteomic studies either from CSF collected after lumbar puncture (Kroksveen et al., 2017) or from a commercial pooled CSF sample (Macron et al., 2018). We found that 34% of proteins detected in the CSF are transmembrane, 21% are cell adhesion molecules, and 10% are both transmembrane and cell adhesion proteins (**Figure 6E, Supplemental Figure S5**). PCDHs are the most prevalent type of transmembrane cell adhesion molecules (**Figure 6F**). Interestingly, although the various subclasses of clustered PCDHs are overrepresented (**Supplemental Figure S5**), non-clustered PCDHs are the most abundant class of proteins in the CSF (**Figure 6G, H**). Altogether, these data and our results strongly suggest that PCDH12 exerts its function through a mechanism involving PCDH12 ectodomain shedding that paracrinally/autocrinally activates downstream signaling such as the WRC.

## Discussion

Like the neurological phenotype in patients lacking PCDH12, the consequence of PCDH12 loss-of-function in our human stem cell -derived cerebral organoid results in a highly deleterious phenotype. Using this model, we were able to pinpoint potential developmental processes involving this protocadherin. Our study is the first to dissect the cellular mechanisms associated with PCDH12 function during cortical development and get some insight into the pathogenic mechanisms involved in *PCDH12*-related NDDs.

Most patients carrying pathogenic PCDH12 variants are comorbid for microcephaly in addition to other neurological conditions. Our findings show that PCDH12-related microcephaly may be caused in part by premature neuronal differentiation, as evidenced by NPCs exiting the cell cycle earlier and differentiating faster than the WT controls (**Figure 3**), resulting in an imbalance between progenitor cell proliferation and differentiation. This cell cycle exit phenotype could potentially be mediated by impaired interaction with centrosomes, as non-clustered δ2 protocadherins have been previously shown to bind with proteins critical for centrosome and spindle function (Robinson et al., 2020). Whether PCDH12 interacts with centrosomal proteins remains to be investigated. Future proximity labeling-based experiments will allow to identify PCDH12 intracellular binding partners in NPCs and neurons.

In addition to its potential role in differentiation, we showed that PCDH12 is unequivocally involved in neuronal migration. *PCDH12*-KO neurons were unable to perform persistent migration, which likely led to abnormal lamination observed in cerebral organoids. Accordingly, PCDH12 patients present with reduction to complete agenesis of the corpus callosum, which is often associated with variants in gene related to neuronal migration (Kato, 2015). In addition, over 70% of PCDH12 patients present with epilepsy (Fazeli *et al*., 2022) for which many known causes involve variants in genes regulating neuronal migration (Guerrini et al., 2008; Guerrini and Parrini, 2010; Reiner et al., 2016).

We also identify a mechanism that implicates recruitment of the WAVE Regulatory Complex at the plasma membrane. Our work agrees with Chen and colleagues who showed that PCDH12 is part of 120 membrane proteins that contain the WRC interacting receptor sequence (WIRS) in their cytoplasmic tail (Chen *et al*., 2014). All non-clustered δ2-PCDHs carry the WIRS, but with various effects on the activity of the WRC: PCDH10, PCHD12, and PCHD19 potentiate it, while PCDH17 inhibits it (Chen *et al*., 2014). This adds another layer of complexity regarding the role of PCDHs during brain development. We speculate that modification of expression of one or another will lead to change in overall actin cytoskeleton dynamics to finely tune cell differentiation and/or migration, in addition to the physiological functions purely associated with their adhesive properties.

Previous studies showed that PCDH12 ADAM10-mediated ectodomain shedding was involved in cell migration (Bouillot *et al*., 2011). The authors hypothesized that in presence of metalloproteinase inhibitors, PCDH12 could not be cleaved and subsequently degraded by the proteasome, increasing cell adhesion that could slow cell invasion of a wounded area. However, we do not think that a mechanism of heightened adhesion is at play here. If increased adhesion was involved, we would expect *PCDH12*-KO neurons to migrate as efficiently as WT neurons in control conditions. Yet, we observed impaired migration with both ADAM10 inhibition in WT and without ADAM10 inhibition in *PCDH12*-KO cells. Our study supports Bouillot and colleagues’ work in how crucial PCDH12 ectodomain shedding by ADAM10 is for exerting its proper biological functions. Interestingly, an ADAM10 conditional KO mouse model targeting progenitor cells also led to premature neuronal differentiation and lamination/neuronal migration defects(Jorissen et al., 2010). Finally, in agreement with our results, recent data have shown that the CSF sheddome is enriched in cell adhesion molecules as well as proteins encoded by NDD risk genes such as neurexins, contactins, or PCDH10 (Martin-de-Saavedra *et al*., 2022).

Altogether, our characterization of *PCDH12*-KO organoids and NPCs provide new insights into the protocadherin function: PCDH12 ectodomain shedding by metalloproteinase ADAM10 plays an important physiological role during brain development, promoting proper cell differentiation and neuronal migration. Further studies are needed to understand how PCDH12 (and generally non-clustered δ2-PCDHs) proteolytic cleavage is determined spatially and temporally, as well as the downstream mechanisms and processes that PCDH ectodomain shedding influences to ensure proper cortical development.

## Acknowledgments

We thank Dr. Talia Lerner for sharing imaging equipment. Live and confocal imaging work was performed at the Northwestern University Center for Advanced Microscopy generously supported by NCI CCSG P30 CA060553 awarded to the Robert H Lurie Comprehensive Cancer Center. This study was supported by the NIH (A.G.-G., grant R00 NS089943) and the Northwestern University Sanger Sequencing Facility.

## Author contributions

AG-G directed the research and generated the CRISPR human stem cell lines. AG-G and JR conceived the experiments and wrote the manuscript.

SLM and DD contributed to immunofluorescence studies. LFT, GY and FF contributed to fractionation and western blot studies. LR, SLM, and ME helped with image analysis. LR analyzed proteomics data. JR analyzed all the data. All authors read and approved the manuscript.

## Methods

### Cell lines

For generating cerebral organoids and neural progenitor cells, feeder-free human male RUES1 embryonic stem cells (hESCs), female GM03651 and male GM03652 induced pluripotent stem cells (iPSCs) were obtained from WiCell and the Coriell Institute, respectively. The cells were not authenticated. *PCDH12*-KO CRISPR lines were generated using the RUES1 cell line. Stem cells were maintained in commercially available mTeSR plus media (StemCell Technologies, #100-0276) at 37ºC and 5% CO_2_. Neural progenitor cells were cultured in DMEM/F-12, HEPES medium (Gibco, #11330032) supplemented with N2, B27, and basic fibroblast growth factor (FGF). Cell culture media was renewed every other day.

### Generation of CRISPR *PCDH12*-KO stem cell lines

RUES1 hESCs were electroporated with px330 (Addgene, #42230) containing a 20-nucleotide single guide sequence targeting exon 1 of human PCDH12. Frame-shifting insertions or deletions, occurring in coding exon 1 of *PCDH12*, were generated in RUES1 hESCs by CRISPR/Cas9 induced non-homologous end joining with primers 5’-CAC CGA GCG CCT CTC TGC GAA CC-3’ and 5’-AAA CGG TTC GCA GAG AGG CGC TC-3’ into BbsI-digested px330. Patient specific c.2511delG variant was also generated by CRISPR/Cas9 induced homologous recombination with primers 5’-ACA CCG TAC AGG ACG CTG CGT AAT CAG-3’ and 5’-AAA ACT GAT TAC GCA GCG TCC TGT ACG-3’ into BbsI-digested px330 and using donor DNA: ACC CTG TAC AGG ACG CTG CGT AAT CAA GGC AAC CAG GGA GCA CCG GCG GAA GCC GAG AGG TGC TGC AAG ACA CGG TCA ACC TCC TTT TCA ACC ATC CCA G. All guides were designed using crispr.mit.edu software. Individual colonies were isolated after 2 weeks, expanded, and genotyped by Sanger sequencing for biallelic mutations. Expression of pluripotency markers was confirmed in all lines and targeted sequencing was used to check for off-target variants.

### Generation of cerebral organoids

Human cerebral organoids were generated from WT and *PCDH12*-KO RUES1 hESCs and WT GM03651 and GM03652 iPSCs with the STEMDiff Cerebral Organoid Kit (StemCell Technologies, 08570) following manufacturer’s instructions. Briefly, embryoid bodies (EBs) were formed from stem cells. After 7 days of culture, EBs were embedded in Matrigel (Corning, 354277) to allow for neuroepithelial expansion. After 10 days of culture, cerebral organoids were maintained using the STEMDiff Cerebral Organoid Maturation Kit (StemCell Technologies, 08571) for up to 40 days.

### Generation of neural progenitor cells

Neural progenitor cells (NPCs) were obtained as previously described (Chambers *et al*., 2009), with stem cells cultured in serum-free floating embryoid body-like aggregates with 1mM of dorsomorphin (Tocris, 3093), 2 mM of A83-01 (Tocris, 2939), and kept shaking at 95 rpm for 7 days. Resultant EBs were plated onto Matrigel-coated (Corning, 354234) dishes in NBF medium made of DMEM/F12 (Gibco, 1330032), 0.5X□N2, 0.5X□B27, and 20 ng/ml of FGF (Invitrogen, PHG0263), to yield rosettes. They were dissociated with Accutase (Millipore, SCR005) after 5 to 7 days. Resultant NPCs were plated onto poly-L-ornithine (PLO)/laminin-coated (Sigma-Aldrich, P3655 and L2020, respectively) dishes in NBF medium. Experiments were performed with NPCs at passages 3 to 8.

### Neurosphere migration assay

Neurosphere migration assay was adapted from previously described protocols (Youn *et al*., 2009). Briefly, NPCs were lifted with Accutase, resuspended in NBF medium, and cultured overnight in suspension in a coating-free dish, shaking at 95 rpm to allow for neurosphere formation. The next day, NBF was replaced with DMEM/F12 supplemented with 1X N2 and B27. Neurospheres were left shaking one more day and subsequently plated onto PLO/Laminin-coated coverslips to trigger radial migration.

Time lapse imaging by phase contrast microscopy was performed 2 hours after plating, for a total of 16 hours using a BioStation IM-Q imaging system (Nikon). Individual cell tracking on the time lapse images was performed using the Manual Tracking plug-in in ImageJ. Directness and Euclidian distance were calculated using the Chemotaxis and Migration Tool software (Ibidi).

### Immunofluorescence

Cerebral organoids were fixed at day 5, 7, 10, 20, and 40 of culture in cold 4% PFA overnight. After three PBS washes, they were transferred to 30% sucrose until sunk, embedded in a gelatin solution and snap-frozen in a dry-ice/ethanol bath. 10-micron thick sections were collected with a cryostat (Leica). Sections were allowed to completely dry before the immunofluorescence process. Then, sections were covered with ∼ 800 μL of 0.1M citrate buffer pH=6 and placed in a steamer (Hamilton Beach, 37350A) for 20 minutes as antigen retrieval treatment. After three washes in a PBS-0.1% Tween20 solution (PBS-T), blocking was performed with 5% Normal Donkey Serum (NDS) in PBS-T for an hour at room temperature. Primary antibodies in 5% Bovine Serum Albumin (BSA) and PBS-T were incubated overnight at 4ºC. The following primary antibodies and concentrations were used: 1:200 N-cadherin (Sigma-Aldrich, SAB5700641), 1:100 phospho-Vimentin (Cell Signaling Technology, 3877S), 1:500 phospho-Histone H3 (Abcam, ab10543); 1:200 Sox2 (Millipore, AB5603), 1:200 Pax6 (Santa Cruz Biotechnology, sc-81649), 1:200 Tbr2 (Abcam, ab23345), 1:200 Satb2 (Abcam, ab51502), 1:500 Ctip2 (Abcam, ab18465), 1:200 Tbr1 (Abcam, ab31940). After three PBS-T washes, species-specific secondary antibodies (Invitrogen, 1:200) diluted in PBS-T were applied for two hours at room temperature, in the dark. Three PBS-T washes were performed, followed by incubation with 2 μ g/mL of Hoechst 33342 (Invitrogen, H3570) for ten minutes, and eventually slides were mounted with ProLong Gold Antifade (Invitrogen, P36930).

For the BrdU experiments, NPCs were plated on an 8-well chambered coverslip (Ibidi, 80821) and treated with a short (30 minutes) or a long (24 hours) pulse of 10 μM BrdU (Sigma-Aldrich, B5002), 24-hour post-plating. At the end of the pulse, NPCs were fixed in cold PFA 4% for 10 minutes, washed three times in PBS, permeabilized 10 minutes in 0.25% Triton X-100, and washed again in PBS. NPCs were then incubated in 2N HCl for 30 minutes and further neutralized with 0.1 M Borate, pH 8.5 for 30 minutes. After three PBS washes, blocking and immunostaining were performed as described below. The following primary antibodies and concentration were used: 1:100 BrdU (BD Biosciences, 347580), 1:100 Ki67 (Abcam, ab15580).

For the TUNEL assay, NPCs were plated on an 8-well chambered coverslip and let to grow for 48 hours. They were then fixed in 4% PFA, permeabilized, and washed in deionized water. The TUNEL staining was performed using the Click-iT Plus TUNEL Assay (ThermoFisher Scientific, C10617) following manufacturer’s instructions.

For the neuronal differentiation experiment, NPCs grown on coverslips were fixed a day after plating, then 7, 14, and 21 days after FGF removal, for 10 minutes in cold 4% PFA. Neurospheres were fixed 2- and 5-days post FGF removal. After three PBS washes, NPCs/neurospheres were permeabilized with 0.5% Triton X-100, and blocked with a 0.3% Triton X-100, 10% NDS, and PBS solution. Primary antibodies in 0.1% Triton X-100, 5% NDS were incubated overnight at 4ºC. The following primary antibodies and concentrations were used: 1:200 Pax6 (BioLegend, 901301), 1:200 Doublecortin (Abcam, ab153668), 1:200 MAP2 (Millipore, MAB3418). After three PBS washes, cells were incubated with species-specific secondary antibodies (Invitrogen, 1:300) diluted in PBS for two hours at room temperature, in the dark. Three PBS washes were performed, followed by incubation with 2 μ g/mL of Hoechst 33342 for ten minutes, and eventually coverslips were mounted with ProLong Gold Antifade.

### Protein extraction, cell fractionation, western blotting

NPCs were grown on a PLO-Laminin-coated 10-cm dish until they reached 70-90% confluency. Membrane proteins were extracted using either 1X RIPA buffer (Cell Signaling Technology, 9806) for whole cell lysates, the Mem-PER Plus Membrane Protein Extraction Kit (Thermo Scientific, 89842), or the Minute™ Plasma Membrane Protein Isolation and Cell Fractionation Kit (Invent Biotechnologies, Inc., SM-005) for fractionated lysates, following manufacturer’s instructions. Briefly, cells were scraped off the plate and centrifuged for 5 minutes at 300 *x g*. After 2 washes, cytosolic proteins were extracted with a permeabilization buffer and centrifugation to isolate them in the supernatant. The remaining pellet was resuspended in a solubilization buffer to isolate membrane proteins. All protein extraction buffers were supplemented with Halt Protease and Phosphatase Inhibitor Cocktail (Thermo Scientific, 78446). Whole cell lysates, cytosolic and membrane fractions were either stored at -80ºC until use, or protein concentration was measured right away using the Pierce BCA Protein Assay Kit (Thermo Scientific, 23225). Protein lysates were denatured in Laemmli buffer by boiling lysates for 5 minutes at 95ºC and directly used for SDS-PAGE or stored at -20ºC until use. Following gel electrophoresis, proteins were transferred to PVDF membranes and then immunoblotted with the following antibodies: cleaved Caspase-3 (1:500, Abcam, ab13847), WAVE1 (1:1000, Millipore, MABN503), alpha-Tubulin (1:1000, Cell Signaling Technology, 2144), N-cadherin (1:1000, Sigma-Aldrich, SAB5700641), cofilin (Abcam, ab11062), and phospho-cofilin (1:1000, Abcam, ab12866). Membranes were then incubated with relevant near-infrared dyes-conjugated secondary antibodies (1:10,000, Li-Cor). Protein signal was detected using the Odyssey® Fc Imaging System (Li-Cor) and quantified with Image Studio (Li-Cor).

### CSF proteomics data analysis

To assess the prevalence of protocadherin species in CSF, we analyzed previously published proteomic studies from samples collected after lumbar puncture (Kroksveen et al., 2017) and from a commercial pooled CSF sample (Macron et al., 2018) in R. The dataset by Macron and coworkers was analyzed by first mapping the accession number to the HGNC symbol and GO term (BiomaRt attribute name_1006). Subsets involved in cell adhesion and/or containing a transmembrane domain were determined using DAVID functional annotation tools. Broad class of each of these 344 transmembrane proteins involved in cell adhesion in CSF was manually determined and then the prevalence of each species was visualized with the ggplot package.

To analyze the protocadherins separately, the dataset generated by Kroksveen and coworkers was subsetted by extracting rows with the “PCDH” pattern in the name. This dataset was merged with a table classifying properties of each protocadherin species using the inner_join function. From this data table, a graph was generated using the ggplot package to show number of unique protocadherin species and/or validated peptide spectra matches in each class and subclass. To look even more specifically at delta-2 protocadherins, the dataset was further subsetted to just this subclass and the number of validated peptide spectrum matches of each gene were visualized using ggplot.

### Statistical analysis

Apart from the proteomics data, all statistical analyses were performed using GraphPad Prism software (v 9.4.1). Shapiro-Wilk or Kolmogorov-Smirnov tests were applied to assess whether assumptions of normality were met. An F-test was applied to assess whether assumptions of equal variance were met. Statistical tests were then chosen accordingly. Numbers (n) of experimental replicates are indicated in the text and the statistical test used is indicated in individual figure legends.

## Supplemental information

**Document S1. Figures S1-S5**

**Movie S1. Live imaging of WT cells migrating out of the neurosphere**.

**Movie S2. Live imaging of *PCDH12*-KO cells migrating out of the neurosphere**.

