## Supplemental document for "Impaired migration and premature differentiation underlie the neurological phenotype associated with PCDH12 loss of function"

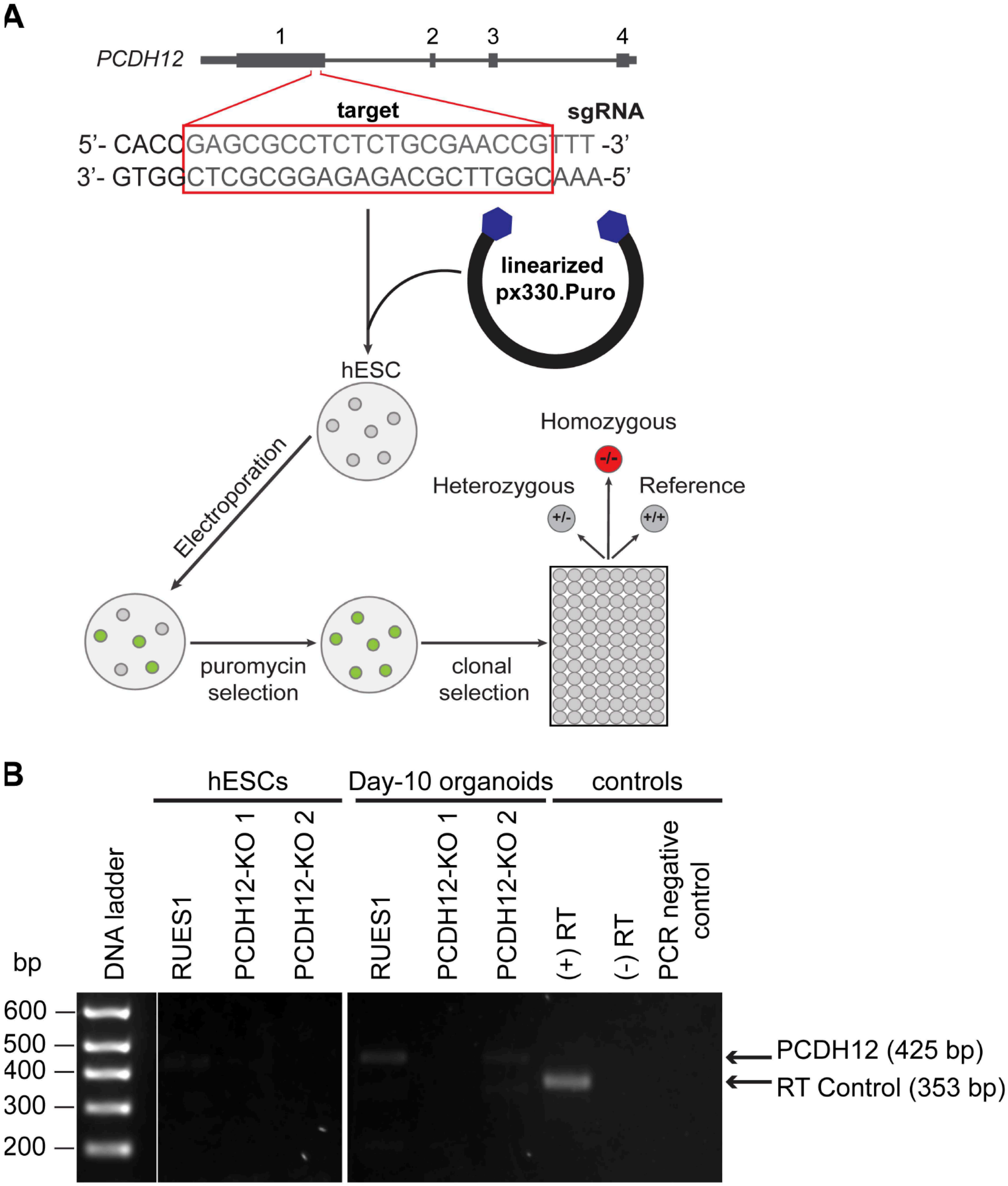


**Supplemental Figure S1. Generation and validation of *PCHD12*-KO stem cell lines, Related to Figure 1.**
(A) CRISPR/Cas9 guides targeting exon 1 are introduced to RUES1 hESCs using an electroporation system. They are then selected for antibiotic resistance.
(B) RUES1 hESC line slightly express PCDH12, and expression increases in 10-day-old cerebral organoids. Expression in PCDH12 CRISPR homozygous cell lines is absent or substantially reduced.

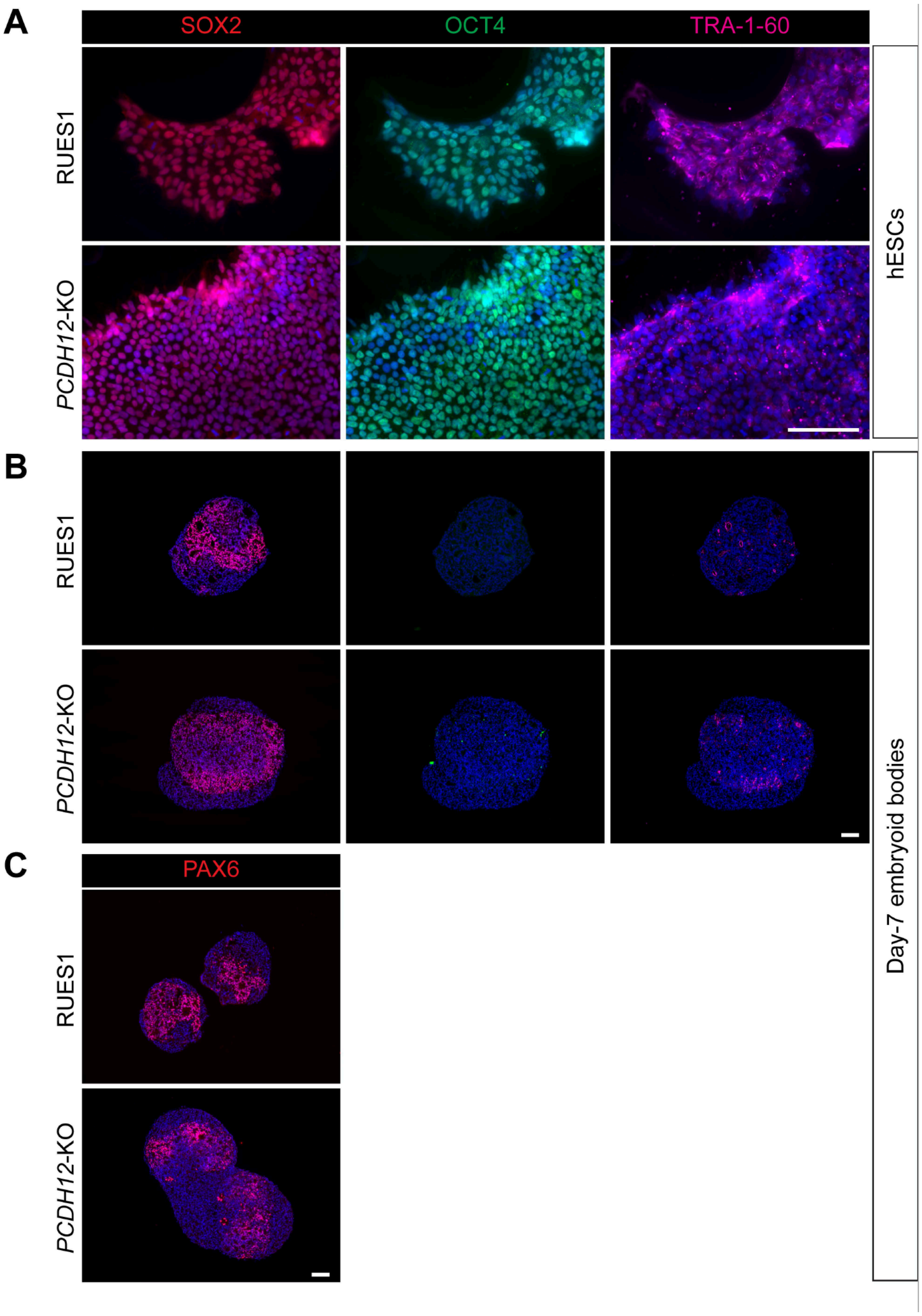


**Supplemental Figure S2. PCDH12-KO stem cells and embryoid bodies express pluripotency markers, Related to Figure 1.**

(A and B) Immunofluorescence images of WT RUES1 (top) and *PCDH12*-KO (bottom) hESCs (**A**) or day-7 embryoid bodies (**B**) stained with SOX2 (red), OCT-4 (green), and TRA-1-60 (magenta). As expected, pluripotency markers' expression decreases as cerebral organoid differentiation progresses.

(C) Immunofluorescence images of WT RUES1 (top) and *PCDH12*-KO day-7 embryoid bodies stained with PAX6 (red) showing specification of progenitors to radial glial cells after a week of differentiation. Scale bars = 100 μm

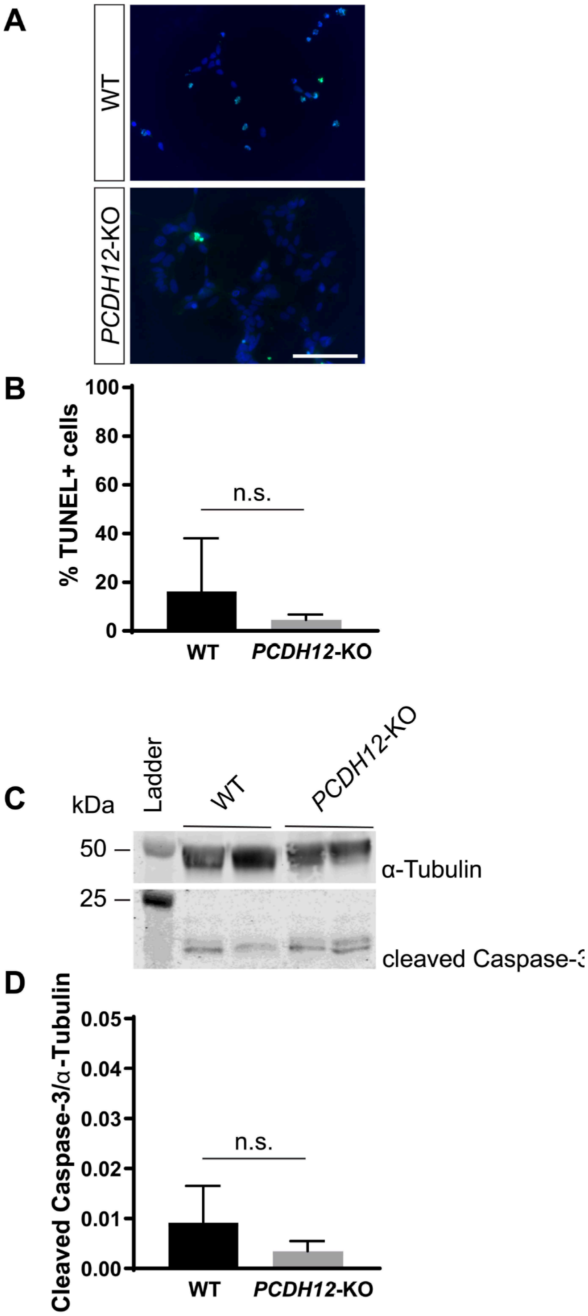


**Supplemental Figure S3. PCDH12 loss of function doesn't affect cell death in NPCs, Related to Figure 3.**

(A) Immunofluorescence images of WT RUES1 (top) and *PCDH12*-KO (bottom) NPCs stained with TUNEL (green).
(B) Quantification of the percentage of TUNEL+ NPCs in WT and PCDH12-KO cultures 48-hours post-plating (n=5 biological replicates for WT; n=5 biological replicates for PCDH12-KO; mean + SD; Mann-Whitney test).
(C and D) Western blot analysis of cleaved-caspase-3 in WT and *PCDH12*-KO NPC whole-cell lysates (n= 7 for WT; n= 7 for *PCDH12*-KO NPCs; mean + SD; Mann-Whitney test).
Scale bar = 100 μm

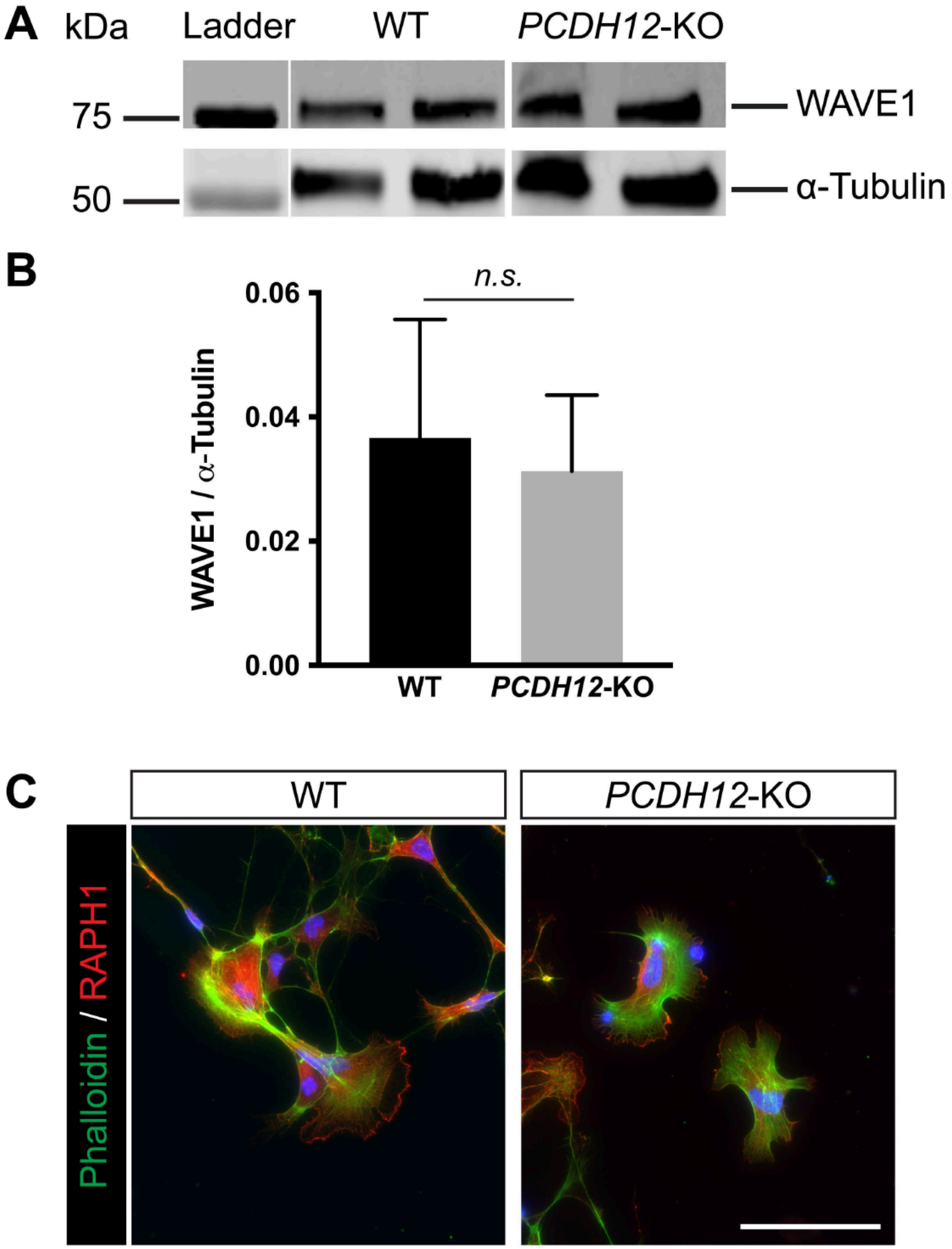


**Supplemental Figure S4. PCDH12 loss doesn't affect overall WAVE1 expression or *PCDH12*-KO neurons' ability to generate lamellipodia, Related to Figure 5.**

(A and B) Western blot analysis of WAVE1 expression in WT and *PCDH12*-KO NPC whole-cell lysates (n= 7 for WT; n= 7 for *PCDH12*-KO NPCs; mean + SD; Unpaired t test).

(C) Immunofluorescence images of WT (left) and *PCDH12*-KO (right) migrating neurons labelled with the lamellipodium marker RAPH1 (red) and F-actin marker phalloidin (green).

Scale bar = 100 μm

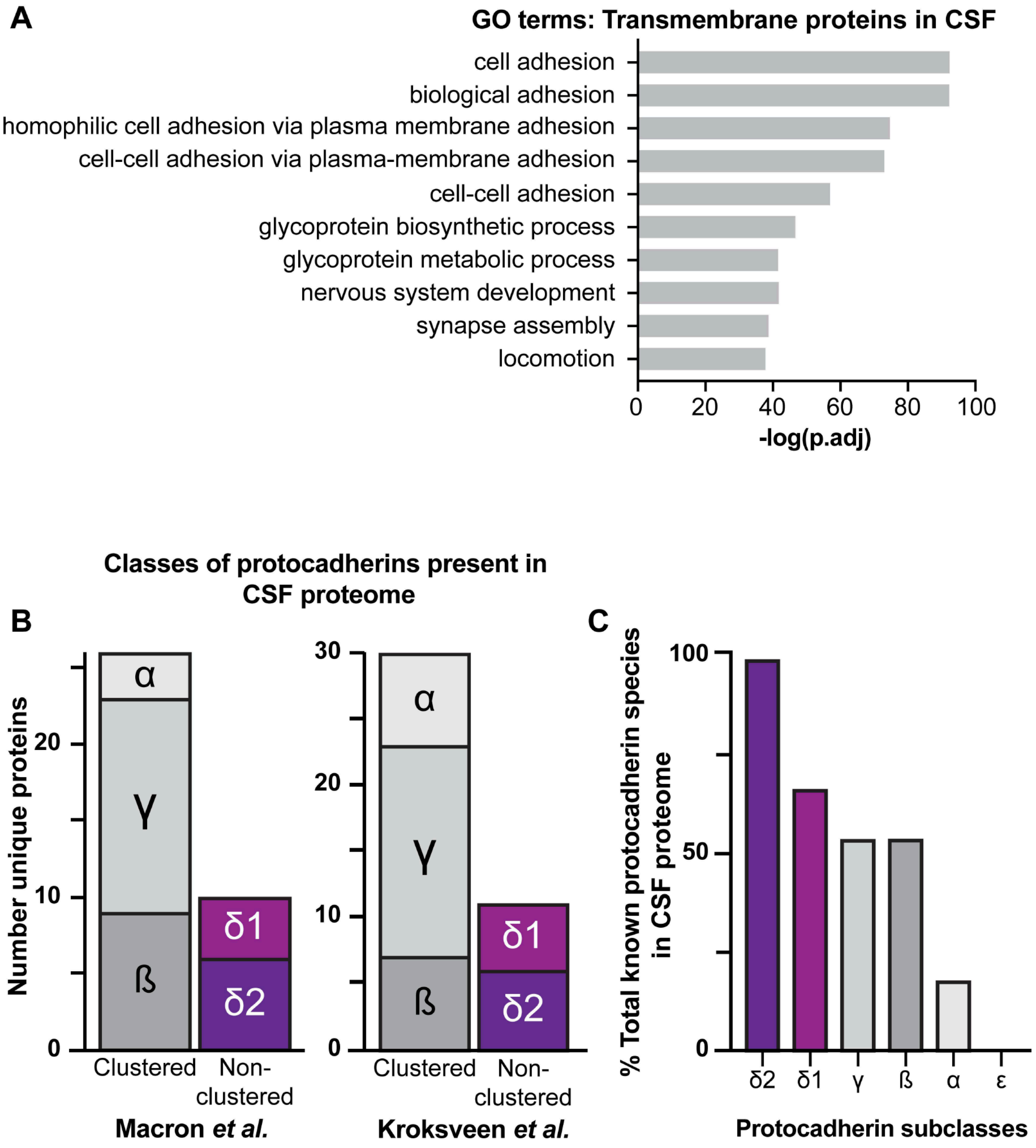


**Supplemental Figure S5. CSF proteome analysis, Related to Figure 6.**
(A) Gene Ontology (GO) term enrichment analysis of the top 10 identified transmembrane proteins in CSF, arranged by fold enrichment.
(B) Quantification of the number of unique protocadherins (PCDHs) detected in the CSF proteome from previously published datasets (Kroksveen et al., 2017; Macron et al., 2018) reveals an overrepresentation of clustered PCDH subgroups.
(C) Histogram showing the relative abundance of protocadherin family members in the CSF proteome.

**Movie S1. Live imaging of WT cells migrating out of the neurosphere.**

**Movie S2. Live imaging of *PCDH12*-KO cells migrating out of the neurosphere.**
